# Development of shuttle vector-based transformation systems for *Chlamydia pecorum* and *Chlamydia caviae*

**DOI:** 10.1101/2024.07.11.603181

**Authors:** Nadja Fässler, Michael Biggel, Martina Jelocnik, Nicole Borel, Hanna Marti

**Affiliations:** Institute of Veterinary Pathology, Vetsuisse-Faculty, University of Zurich, Zurich, 8057, Switzerland; Institute of Food Safety and Hygiene, Vetsuisse-Faculty, University of Zurich, Zurich, 8057, Switzerland; School of Science, Technology and Engineering, University of the Sunshine Coast, Sippy Downs, 4556, Queensland, Australia; Center for Clinical Studies, Vetsuisse-Faculty, University of Zurich, Zurich, 8057, Switzerland

**Keywords:** *Chlamydiaceae*, *Chlamydia*, plasmid, horizontal gene transfer, zoonoses

## Abstract

*Chlamydia (C.) abortus*, *C. caviae* and *C. pecorum* are obligate intracellular, zoonotic pathogens, which have all been associated with community-acquired pneumonia in humans. *C. abortus* is the causative agent of enzootic ovine abortion in small ruminants and can lead to miscarriage in women. *C. caviae* causes conjunctivitis in guinea pigs, while *C. pecorum* is found in livestock, resulting in economic losses and contributing to the decline of the koala population in Australia. Studying the biology of these bacteria has been challenging due to a dearth of genetic tools. This study aimed to establish transformation systems for *C. abortus* and *C. pecorum* using shuttle vectors and to expand upon already existing protocols for *C. caviae*. Shuttle vectors comprised the cryptic plasmid of the chlamydial species of interest, the pUC19 origin of replication (*ori*), a beta-lactamase (*bla*), and genes that mediate heterologous expression of fluorescent proteins (GFP, mNeonGreen, mScarlet). A *C. suis*-tailored transformation protocol and a previously established protocol for *C. psittaci, C. trachomatis* and *C. pneumoniae* were applied. While *C. pecorum* and *C. caviae* transformation experiments were successful, transformation of *C. abortus* remained ineffective. Shuttle vectors yielded stable transformants over several passages in the presence and absence of selective antibiotics while the fluorescence intensity of GFP was superior compared to mNeonGreen. Finally, we co-cultured GFP- and mScarlet-expressing *C. pecorum* strains demonstrating that both fluorophores can be detected in the same cell or even inclusion, possibly promoting homologous recombination. These findings open new avenues into our understanding of interstrain and interspecies co-infection dynamics both *in vitro* and *in vivo*.

## Introduction

Several chlamydial species of the obligate intracellular bacteria family *Chlamydiaceae,* found in livestock, pets, and wild animals, harbor a zoonotic potential (1). In their animal hosts, severity of the disease ranges from asymptomatic to mild conjunctivitis to life-threatening pneumonia and abortion, depending on the pathogen and the affected host (1,2). A well-known zoonotic *Chlamydia* species is *C. abortus*, which harbors two different strain groups, of which one is responsible for enzootic abortion in small ruminants and the other has been described in birds (3,4). Both strain groups may lead to community-acquired pneumonia (CAP) in humans (1,5,6) and ruminants strains have been shown to induce miscarriage in pregnant women (7). Other veterinary chlamydial species have also been reported to cause CAP, such as *C. caviae*, a conjunctivitis agent in guinea pigs (8). *C. pecorum* possesses a similar zoonotic potential (9). However, it is primarily considered a veterinary pathogen with a wide array of animal hosts ranging from livestock to marsupials. In the koala, *C. pecorum* causes debilitating diseases such as blindness and infertility and is a major cause of the population decline in this protected species (1,10). In livestock, the clinical impact of *C. pecorum* infections depends on the animal host and the geographical region: While strains causing polyarthritis and abortion in ruminants, as well as encephalomyelitis in calves, have mostly been reported in Australia and New Zealand, the economic and clinical impact of *C. pecorum* on livestock in Western Europe remains understudied (1,10).

Studies into genetic diversity have shown that *C. pecorum* is phylogenetically diverse and variable in terms of strain-dependent virulence (11–16) comprising virulence factors such as coding tandem repeats in *incA* and the ORF663 loci (16). Moreover, deletions in the native plasmid could potentially be responsible for the virulence of abortigenic *C. pecorum* sequence type 23 (12). The genetic diversity of *C. caviae* has only recently been investigated revealing the circulation of few but distinct strains within the European guinea pig population (17,18). One *in vitro* study identified *incA* and SinC as potential virulence factors (19). In contrast, ruminant *C. abortus* strains are highly conserved and have not revealed any distinct virulence groups (20). Overall, in-depth analysis of proposed virulence factors of these zoonotic veterinary chlamydiae have been hampered by a dearth of genetic tools. While such tools have recently been developed for the *Chlamydia* genus as a whole, most studies have focused on human chlamydiae, such as *C. trachomatis* and *C. pneumoniae*, as well as the animal model species *C. muridarum* (21,22), except for individual studies investigating *C. caviae* (19), *C. felis* (22)*, C. psittaci* (23), and *C. suis* (24).

This slow development of genetic tools has also hindered our understanding of chlamydial co-infection dynamics, an important but understudied field in chlamydial biology. The importance of co-infections *in vivo* is highlighted by many reports about the presence of different chlamydial species and strains in the same animal or human host (20,25–28). While various *in vitro* co-culture protocols exist (29–32), the study of such co-infection dynamics have so far been limited to species-level analysis using antibodies (30,32).

To fill these gaps, we established a calcium-chloride (CaCl_2_)-mediated, shuttle vector-based transformation system for *C. pecorum* by which we gained insights into intraspecies co-infection dynamics between different *C. pecorum* livestock and koala strains. Furthermore, we explored the fate of the native plasmid during transformation, attempted transformation of *C. abortus,* and expanded upon existing transformation protocols for *C. caviae*.

## Material and Methods

### Media and cell culture

Experiments were performed with LLC-MK2 cells (rhesus monkey kidney cell line, kindly provided by IZSLER, Brescia, Italy), cultured at 37 °C with 5 % CO_2_ as previously described (33) with few modifications (24). Briefly, we used antibiotic-free growth media consisting of minimal essential medium with Earle’s salts, 25 mM HEPES, L-Glutamine-free (GIBCO, Invitrogen, Carlsbad, CA, USA), 10% fetal calf serum (FCS, BioConcept, Allschwil, Switzerland), 2 mM GlutaMAX (200 mM, GIBCO), and 0.4 g D-(+)-glucose (Sigma-Aldrich, St. Louis, MO, USA). Infection medium contained 5-fold higher glucose concentrations and was supplemented with cycloheximide (1.5 µg/mL, Sigma-Aldrich), if needed. Sucrose phosphate glutamate (SPG) buffer with 218 mM sucrose (Sigma-Aldrich), 3.76 mM KH_2_PO_4_ (Sigma-Aldrich), 7.1 mM K_2_HPO_4,_ and 5 mM GlutaMAX (GIBCO) was used for long-term storage of *Chlamydia* stocks as described (34).

### *Chlamydia* stocks

For this study, the *C. pecorum* strains were generously provided by the Centre for Bioinnovation, University of the Sunshine Coast, Sippy Downs, Australia (strains MC/MarsBar_2018 and IPA), by the Istituto Zooprofilattico Sperimentale della Lombardia e dell’Emilia Romagna, IZSLER, Italy (strain PV7855) or were generous gifts from J. Storz, Baton Rouge, LA, USA (strains W73, P787, 1710S). The *C. caviae* strain was kindly provided by Dr. Barbara Sixt, Umea University, Sweden, and originated from Roger Rank (University of Arkansas for Medical Sciences). The avian *C. abortus* strain 15-70d24 was kindly provided by Dr. Monika Szymanska-Czerwinska, Department of Cattle and Sheep Diseases, National Veterinary Research Institute, Pulawy, Poland, and the ruminant *C. abortus* strain was a gift from Dr. Jones (Moredun Research Institute, Edinburgh, UK) (Table 1).

**Table 1:** List of strains used in this study.

| Species | Strain | Host | Disease | Plasmid | Reference |
| --- | --- | --- | --- | --- | --- |
| *C. pecorum* | P787 | Sheep | Polyarthrititis | No | (35,36) |
| *C. pecorum* | MC/MarsBar_2018 | Koala | Chronic cystitis | Yes | (37,38) |
| *C. pecorum* | IPA | Sheep | Polyarthrititis | Yes | (39) |
| *C. pecorum* | PV7855 | Chamois | Pneumonia | Yes | (35) |
| *C. pecorum* | 1710S | Pig | Inapparent intestinal infection | No | (39) |
| *C. caviae* | GPIC | Guinea pig | Conjunctivitis | Yes | (19) |
| *C. abortus* | S26/3 | Sheep | Abortion | No | (40) |
| *C. abortus* | 15-70d24 | Avian | No information | Yes | (41) |

Stocks of semi-purified *Chlamydia* were prepared according to established protocols (31). Briefly, chlamydiae were grown for up to five passages, aiming for 75 – 100 % infection in T75 flasks (TPP, Trasadingen, Switzerland). Destruction of infected cells was achieved by scraping them into the supernatant and mechanical disruption through vigorous vortexing with sterile glass beads (5 mm diameter, Sigma-Aldrich) for 1 min. Cellular debris was removed by centrifugation (500 *g*, 4 °C, 10 min) and the chlamydiae pelleted with 10’000 *g* at 4 °C for 45 min. Resuspension was performed in SPG, aiming for a stock concentration ranging from 10^9^ to 10^10^ inclusion forming units per mL (IFU/ml). Aliquots were then stored at -80 °C until use.

The exact stock concentration was expressed as IFU/ml and determined by subpassage titration performed in duplicate as described (24). In brief, SPG stocks were inoculated onto 96-well plates (TPP) followed by a threefold serial dilution. After centrifugation (1’000 *g*, 1 h, 25 °C), inocula were replaced with infection medium containing cycloheximide and incubated at 37 °C for 48 h. Monolayers were then fixed in ice-cold methanol for 10 min, and an immunofluorescence assay (IFA) was performed to visualize chlamydial inclusions using the protocol described by Leonard et al. (42). In brief, cells were blocked for 30 min in phosphate-buffered saline (PBS, GIBCO) supplemented with 1 % bovine serum albumin (BSA, Sigma-Aldrich) after methanol-fixation. Chlamydial inclusions were then labeled with a *Chlamydiaceae*-specific primary antibody diluted 1:200 in blocking solution (*Chlamydiaceae* LPS, Clone ACI-P; Progen, Germany). After a washing step in PBS, DNA was stained with 1 μg/mL 4’, 6-diamidino-2’-phenylindole dihydrochloride (DAPI, Molecular Probes, Eugene, OR, USA) and inclusions were visualized for 45 min with the secondary antibody Alexa Fluor 488 goat anti-mouse (Molecular Probes) diluted in blocking solution (1:500). Inclusion numbers was then counted in a whole well using the Nikon Ti Eclipse epifluorescence microscope (Nikon, Tokyo, Japan), which was used to determine IFU/ml.

### Shuttle vector construction and vector stock preparation

The detailed protocol is published and was first described in a previous study (24) (https://dx.doi.org/10.17504/protocols.io.kxygxy53wl8j/v1; Chapters 1-3). Briefly, *Chlamydia* shuttle vectors were designed to comprise an origin of replication (*ori*) of *E. coli* (pUC) together with a fluorescent gene (mNeonGreen, mScarlet, GFP), an ampicillin resistance gene (*bla*), and the entire native plasmid sequence of the chlamydial species of interest. For *C. pecorum*, the native plasmid sequence was obtained from strain W73 (40,43). For *C. abortus* and *C. caviae*, avian strain 15-70d24 and strain GPIC were used, respectively (Table 1). Primers for all shuttle vectors, the exact composition, and vectors used for shuttle construction are listed in Table S1. Vector construction was performed by PCR amplification with overlapping primers using Phusion Hot Start II DNA Polymerase kit (ThermoFisher Scientific, Waltham, MA, USA) and assembly was performed with the HiFi DNA Assembly Cloning Kit (New England Biolabs, NEB; Ipswich, MA, USA). Assembled vectors were then transformed into NEB5alpha *E. coli*. Vector sequences were confirmed by whole-plasmid sequencing using the Full PlasmidSeq service based on long-read sequencing technology performed by Microsynth AG (Balgach, Switzerland). Unmethylated vector DNA for chlamydial transformation was then produced in *dam-/dcm-E. coli* (NEB) as recommended (44). Vectors and fluorescent genes used for construction included the following plasmids: pL0M-S-mNeonGreen-EC18153, a gift from Julian Hibberd (Addgene plasmid #137075) (45), pmScarlet_C1, a gift from Dorus Gadella (Addgene plasmid # 85042) (46), and pBOMB4R, kindly provided by Ted Hackstadt, MT, USA (47).

### Transformation of *Chlamydia*

Transformation was attempted with two different protocols. Both protocols are published (https://dx.doi.org/10.17504/protocols.io.kxygxy53wl8j/v1, Chapter 4). **Protocol A** was optimized for *C. suis* (24). Briefly, 6.25 x 10^6^ IFU of *Chlamydia* was mixed with 7.5 µg of shuttle vector in 100 µl of calcium chloride (CaCl_2_) solution containing 100 mM CaCl_2_ (Merck) and 20 mM Tris (Roche, Basel, Switzerland), with a pH adjusted to 7.4. Incubation was performed at room temperature for 1 h with mild agitation, followed by the addition of freshly trypsinized cells resuspended in 100 mM CaCl_2_ and seeding onto 6-well plates. After centrifugation (1’000 *g*, 35 °C, 1 h), cultures were incubated for 6 h before addition of 1.5 µg/ml cycloheximide and 0.5 µg/ml ampicillin (ThermoScientific). Up to four passages were performed every 36-96 h, with possible positive cultures appearing after two passages. For the collection of successful transformants, samples were scraped into SPG and frozen at -80 °C until further processing. A fresh coverslip seeded to confluence in a 24-well plate was infected with a 1:100 to 1:200 dilution of the culture for imaging of positive cultures. After centrifugation (1’000 *g*, 25 °C, 1 h) and an incubation period of 48 h, cultures were fixed with ice-cold methanol for 10 min. Coverslips were washed with PBS, stained with DAPI (1 μg/mL) for 10 min, and transferred to glass slides with FluoreGuard mounting medium (Hard Set; ScyTek Laboratories Inc., Logan, UT, USA). Imaging was performed as described in the section “Fluorescence Microscopy and Imaging.”

If three attempts with this protocol remained unsuccessful, transformation was attempted with **Protocol B**, as previously described (22,23,48), with few modifications. Briefly, 6.25x10^6^ IFU of *Chlamydia* mixed with 15 µg of shuttle vector were incubated for 30 min at room temperature in 50 mM instead of 100 mM CaCl_2_. Cells were trypsinized and suspended in the same concentration of CaCl_2_. These suspensions were then mixed together and incubated for another 20 min at room temperature before they were processed as described above.

### Recovery of clonal stocks

Clonal purification was performed with a protocol derived from published work (44). Briefly, a 96-well plate (TPP) was divided into four quadrants. Two quadrants per plate were used per strain. The plate was seeded to confluence with LLC-MK2 cells, and the first well of a quadrant was inoculated with 10 µl of undiluted transformant stock. The first well of the second quadrant was inoculated with 1:100 diluted transformant stock. In both quadrants, a three-fold serial dilution was first performed vertically in the first column, followed by horizontal serial dilution, resulting in 24 individually diluted wells per quadrant and 48 wells in total. After centrifugation (1’000 *g,* 33 °C, 1 h), inocula were replaced with infection medium supplemented with 5 µg/ml ampicillin. After an incubation period of 48 h at 37 °C, 96-well plates were investigated with a Nikon Ti Eclipse epifluorescence microscope (Nikon) and individual wells were selected that contained 0-1 visible fluorescent inclusions. These wells were scraped and passaged two times in 24-well plates (TPP) and one time in 6-well plates (TPP) in selective antibiotics aiming for only few inclusions, an infection rate of 50-90% and 75-100% in Passages 1, 2 and 3, respectively. Stocks were then scraped into SPG, stored and their concentration determined as described above. The detailed protocol is published (https://dx.doi.org/10.17504/protocols.io.kxygxy53wl8j/v1, Chapter 6).

### Stability Assay

Clonally purified stocks were inoculated onto 13 mm glass coverslips (ThermoFisher Scientific) in 24-well plates seeded with LLC-MK2 cells aiming for a multiplicity of infection (MOI) of 0.1 and passaged five times every 48-72 h by scraping the wells and using 1:500 of the culture to infect fresh cells, both in the presence and absence of 5 µg/ml ampicillin. After initial infection (P0) and at the fifth passage (P5), coverslips were fixed in methanol, stained with DAPI, and mounted as described above in the section “Transformation of *Chlamydia.*” Two coverslips per condition (with or without ampicillin) were semi-quantitatively assessed by counting 100 inclusions either at 400x or 1’000x magnification with a Leica DMLB fluorescence microscope. Each inclusion was identified by DAPI and the presence or absence of the appropriate fluorophore. Next, the number of fluorescent inclusions was expressed as percentage of total inclusions.

### Fluorescence Microscopy and Imaging

All images were taken with a Nikon Upright Microscope Eclipse N*i*-U using either a 40X (Nikon Plan Apo 40x/0.95, OFN25 DIC N2) or 60X (Nikon Plan Apo 60x/0.95, OFN25 DIC N2) objective, and a Nikon DS Ri2 camera. For image capture, the manual multichannel capture function of the NIS-Elements AR software (v. 4.30.02, 64-bit) was used. DAPI channel, FITC channel (GFP, mNeonGreen), and TRITC channel (mScarlet) exposure times were 10-30 ms, 2 sec, and 5 s, respectively. The mean fluorescence intensity per inclusion was measured with the area analysis function by autodetecting the area of 30 individual inclusions per strain of interest.

### Statistics

GraphPad Prism (v. 10, GraphPad Software, Boston, MA, USA) was used for all statistical analyses. The Mann-Whitney test was used to compare two means of data that were not normally distributed, and an unpaired t-test was used if the data was normally distributed.

### Whole-genome sequencing and analysis

Genomic DNA was extracted using the Qiagen Blood&Tissue kit (Qiagen, Hilden, Germany) according to manufacturer’s instructions. Strain identity of transformed cultures was first confirmed by sequencing of the major outer membrane protein (*ompA*) with Sanger sequencing at Microsynth AG. For *C. pecorum*, a species-specific full-length *ompA* PCR was used (39) as previously described (15). For *C. abortus* and *C. caviae*, an in-house PCR with the more general *ompA* primer pair BGP4 (5′-ATGAAAAAACTCTTGAAATCGG-3′) and CT90UF (5′-ACTGTAACTGCGTATTTGTCTG-3′) (49) was applied. The detailed protocol is published (https://dx.doi.org/10.17504/protocols.io.kxygxy53wl8j/v1, Chapter 5).

For long-read sequencing, libraries were prepared using the RBK114-24 Rapid Barcoding Kit and sequenced on a MinION device using R10.1.4 flow cells (Oxford Nanopore Technologies). Basecalling, adapter trimming, and demultiplexing were performed with Dorado (basecall server 7.3.9, SUP mode) implemented in MinKNOW 24.02.6 (Oxford Nanopore Technologies). Host sequences were removed by mapping reads to the *Macaca mulatta* genome (Accession No. GCF_003339765.1) and collecting unmapped reads using minimap2 v2.24 (50) and samtools 1.15.1 (51). Following read filtering to a minimum length of 1000 bp with nanoq v0.10.0 (52), assemblies were generated with Flye v2.9.3 (53).

Genome assemblies are available under NCBI Bioproject No. PRJNA962280.

## Results and Discussion

### *C. pecorum* is stably transformed with 100 mM CaCl_2_ for 1 h at room temperature

Using transformation Protocol A, optimized for *C. suis* (24), we attempted transformation of *C. pecorum* strains with a shuttle vector termed pUC-Cpecpl-GFP, comprising the whole-plasmid sequence of *C. pecorum* strain W73 and expressing ampicillin resistance as well as a GFP fluorescence protein (Figure 1A, Table S1). It was derived from pBOMB4R by Bauler and Hackstadt (47).

**Figure 1.**
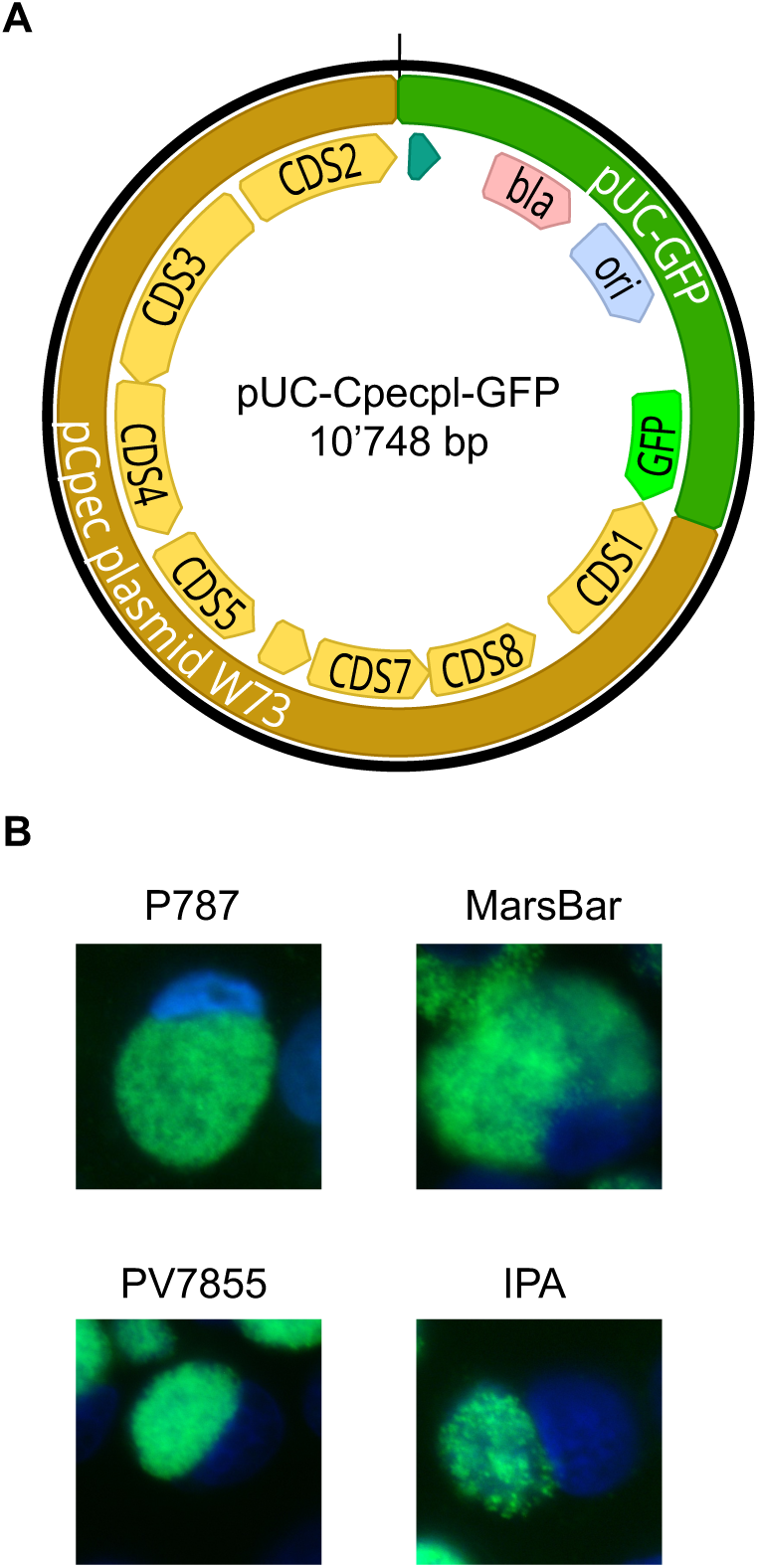
Transformation of *C. pecorum* strains with shuttle vector pUC-Cpecpl-GFP. **A)** Shown is shuttle vector pUC-Cpecpl-GFP comprising the whole plasmid of *C. pecorum* strain W73 (labeled in light brown; coding sequences CDS 1-8 labeled in yellow) and pUC-GFP consisting of a beta-lactamase (*bla,* pale red), the pUC origin of replication (*ori*, pale blue) and GFP (neon-green). **B)** Shown are representative images of *C. pecorum* strains P787, MC/MarsBar_2018, PV7855, and IPA successfully transformed with pUC-Cpecpl-GFP at the time of collection after 3-4 passages. Images for DAPI (blue) and GFP (green, FITC channel) were taken individually, using a 60x objective, and merged.

Using *C. pecorum* strains P787, MC/MarsBar_2018, PV7855, IPA and 1710S, we achieved successful transformation with all strains (Table 1, Figure 1B) except for the porcine strain 1710S (image not shown). First transformants were observed after the second passage, similar to *C. suis* (24). Unsuccessful transformation attempts were repeated three times. Next, to ensure the identity of each strain before further processing, the near full-length *ompA* sequence was obtained by Sanger sequencing and compared against the NCBI databank with the Nucleotide Basic Local Alignment Search Tool (BLASTn) (https://blast.ncbi.nlm.nih.gov/Blast.cgi) (Table S2A). As a next step, confirmed strains were clonally purified, and the stability of the shuttle vector was tested in the presence and absence of ampicillin over five passages (Table S2B). In 200 counted inclusions per strain and condition, we detected no more than two GFP-negative inclusions. This number was independent of the passage number or the presence/absence of selective antibiotics (Table S2B). These results suggest that all transformants stably maintained the pUC-Cpecpl-GFP shuttle vector, including strain P787, a strain that naturally lacks a native chlamydial plasmid (35,36). These findings coincide with what other studies have found for plasmid-free chlamydial strains of *C. trachomatis* (54), *C. muridarum* (55) and *C. pneumoniae* (22). The overall stability of the *C. pecorum* transformants was expected, considering that shuttle vectors containing the complete plasmid sequence of the targeted chlamydial species have yielded stable transformants for many different chlamydial species, including *C. trachomatis* (48), *C. muridarum* (55), *C. suis* (24), *C. pneumoniae* (22) and *C. psittaci* (23). Plasmid stability is likely conferred by the genes responsible for plasmid maintenance (56), namely *pgp* 1, 2, 6 and 8 corresponding to coding sequences (CDS) 3, 4, 8 and 2, respectively (55). Moreover, the stabilizing effect of *pgp* 6 was later used to develop suicide vectors for *C. trachomatis* by placing the gene under tetracycline regulation (44). The importance of the CDS for plasmid stability was further demonstrated by the development of vectors that contained only the chlamydial *ori* (57): the vectors were quickly lost within the first three passages in the absence of antibiotic selection. However, stability may also depend on the vector backbone used as well as the chlamydial species of interest. For example, another study found that a broad-spectrum plasmid from *Bordetella pertussis* yielded stable transformants in *C. trachomatis,* with the vector being maintained as both plasmid and episome (58).

Moreover, one of our previous studies showed that *C. suis* only requires sufficient homologous chromosomal regions but not the chlamydial plasmid for stable transformation and allele replacement (24). In the present study, we did not further investigate the specific factors necessary for vector maintenance in *C. pecorum.* Transformation of *C. pecorum* was attempted in another study using Protocol B with an incubation period of 30 min in 50 mM CaCl_2_ at room temperature, followed by 20 min of incubation with trypsinized cells before centrifugation and incubation (22). In that study, a shuttle vector designed for *C. pneumoniae* was used, but attempts to transform *C. pecorum* remained unsuccessful (22), possibly due to species barriers based on CDS 2 (*pgp 8*) as found for other chlamydial species (59). We also used Protocol B for the transformation of strain 1710S using pUC-Cpecpl-GFP. However, these attempts remained unsuccessful (data not shown), indicating that Protocol B does not yield a higher transformation rate for *C. pecorum* than Protocol A.

### *C. caviae* was transformed with Protocol B but not Protocol A

While initial trials failed to transform *C. caviae* with Protocol A, we used both Protocol A and Protocol B in parallel, with one minor change due to the high infectivity of this chlamydial species: Instead of 0.5 µg/ml ampicillin for selection at 6 hours post infection, we used 5 µg/ml. We were able to introduce shuttle vector pUC-Ccavpl-GFP into *C. caviae* strain GPIC with 30 min in 50 mM CaCl_2_ at room temperature, followed by co-incubation with trypsinized cells for 20 min (Protocol B) after only one attempt (Figure 2). The number of transformed inclusions after the second passage was higher than *C. pecorum*. Interestingly, transformation attempts with Protocol A as well as an alternative protocol with a 30-min incubation in 100 mM CaCl_2_ followed by cell co-incubation for 20 min remained unsuccessful (Table 2). These findings demonstrate that an increase in the CaCl_2_ concentration does not necessarily increase transformation efficiency. Furthermore, they are indicative for a complex interplay between individual factors that contribute to successful transformation.

**Figure 2.**
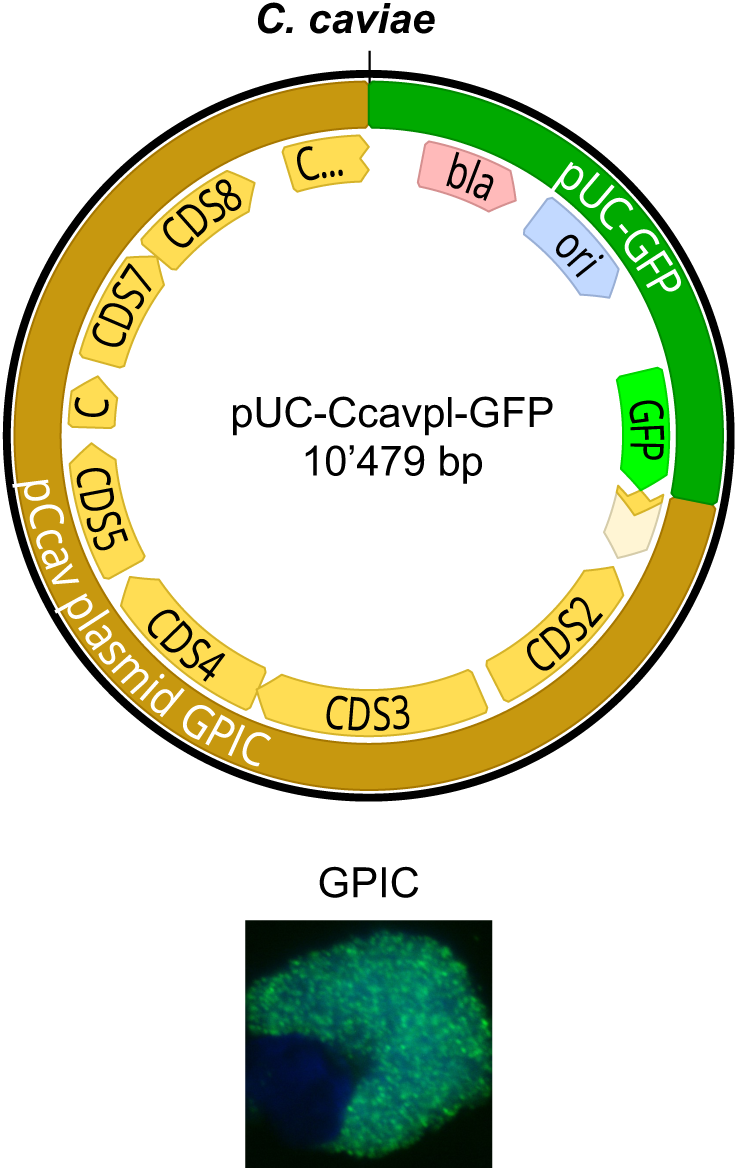
Transformation of *C. caviae* with shuttle vector pUC-Ccavpl-GFP. Shown is shuttle vector pUC-Ccavpl-GFP comprising the whole plasmid of *C. caviae* strain GPIC (labeled in light brown; coding sequences CDS 1-8 labeled in yellow). The shuttle vector possesses pUC-GFP which contains a beta-lactamase (*bla,* pale red), the pUC origin of replication (*ori*, pale blue) and GFP (neon-green). The bottom of the panel shows a representative image of successfully transformed *C. caviae* GPIC. Images for DAPI (blue) and GFP (green, FITC channel) were taken individually, using a 60x objective, and merged.

**Table 2:** Overview of transformation protocols used in this study.

| ***Chlamydia* species** | ***C. pecorum*** | ***C. caviae*** | ***C. abortus*** |
| --- | --- | --- | --- |
| **Protocol A**<br>[100 mM, 1h] | Yes | No | No |
| **Protocol B**<br>[50 mM, 30 min + 20 min] | Not attempted | Yes | No |
| **Alternative Protocol**<br>[100 mM, 30 min + 20 min] | Not attempted | No | No |

With the successful transformation of *C. caviae* using a shuttle vector, we expand upon the existing genetic toolbox of this species (19). This GFP-expressing strain could be used in guinea pig infection models to better understand the dissemination of chlamydiae in an infected host over time. In the past, guinea pig models have been used to mimic genital *C. trachomatis* infections (60) but have also been used as a conjunctivitis model to investigate trachoma-like infections (61).

These protocols were also used in an attempt to transform a ruminant (S26/3) and an avian (15-70d24) *C. abortus* strain using a shuttle vector that contains the whole-plasmid sequence of strain 15-70d24 (Figure S1). However, all attempts to transform these strains remained unsuccessful (Table 2). Whole-genome analyses of ruminant *C. abortus* have revealed that these strains fall into a highly conserved clade and form a distinct cluster that does not undergo homologous recombination (20). This genomic inertia could explain the lack of transformation success, especially considering that ruminant *C. abortus* strains do not carry a natural plasmid in contrast to the recently characterized avian strains (4,62,63). Although it has been speculated that the plasmid could be used for *C. abortus* transformation (63), this present study demonstrates that further modifications of existing protocols are necessary to genetically tag *C. abortus*.

### If present, the native plasmid is replaced by the shuttle vector upon transformation

All transformants were confirmed by Nanopore whole genome sequencing to ensure the identity of all transformed strains generated in this study (Supplementary Table S3). In addition, we screened for the presence of a native plasmid to confirm that the shuttle vector entirely replaces the native plasmid, as has been observed previously (24,48). Our results confirm the strain identity of all transformants and further show that, when present in the original strain, the native plasmid is entirely replaced by the shuttle vector, as expected. Interestingly, sequencing of the *C. caviae* transformant revealed two contigs for the plasmid despite high coverage. To ensure the identity of the vector sequence of our *C. caviae* transformant, we transformed the shuttle vector back into *E. coli* and performed full-plasmid sequencing as described previously (24). Using this method, we confirmed that the vector used for transformation was identical to the vector that could be retrieved from GFP-expressing *C. caviae* (Supplementary Table S3).

### GFP-tagged inclusions yield higher fluorescence intensities than mNeonGreen-tagged chlamydiae and may be used for co-infections with mScarlet-positive chlamydiae

Culture-independent sequencing methods of clinical samples have shown that more than one *C. pecorum* strain may infect a single host (27). These intriguing findings suggest complex co-infection dynamics of different chlamydial strains within the same host and possibly within the same host cell. Heterologous expression of different fluorophores could be a helpful tool for exploring such co-infection dynamics *in vitro*. GFP has an excitation maximum (λ_ex_) of 488 nm and an emission maximum (λ_em_) of 510 nm (https://www.fpbase.org/). Consequently, the expected bleed-over between GFP and mScarlet (λ_ex_/λ_em_ = 569/594 nm) is expected to be limited. For co-culture experiments, we generated a *C. pecorum* shuttle vector termed pUC-Cpecpl-Scarlet in which GFP was replaced with mScarlet (Table S1). Furthermore, we designed a shuttle vector with mNeonGreen (pUC-Cpecpl-NeonGreen; λ_ex_/λ_em_ = 506/517 nm) because the fluorophore is expected to be brighter and less sensitive to photobleaching than GFP (64). We used *C. pecorum* strains P787, MC/MarsBar_2018, IPA, and 1710S for transformation and were able to transform all strains with pUC-Cpecpl-NeonGreen, and all strains except for 1710S with pUC-Cpecpl-Scarlet (Figure 3A). The mean fluorescence intensity of 1710S tagged with mNeonGreen was significantly lower than other mNeonGreen-positive strains (Figure S2). Taken together with the overall challenge to successfully transform 1710S, and considering the fact that 1710S does not naturally carry a plasmid (39), factors other than the plasmid sequence may play a role in plasmid maintenance of this strain and possibly other chlamydiae such as *C. abortus*. The weak fluorescence could indicate a lower number of plasmid copies in the transformed 1710S compared to other strains. However, since plasmid copy numbers in chlamydiae depend on various factors including the chlamydial developmental cycle (61), comparing interstrain and interspecies differences remains difficult. Suboptimal growth conditions for 1710S could also affect their low transformation rate. A previous study looking into *in vitro* growth characteristics between different *Chlamydia* strains, including 1710S, showed that CaCo2 cells yield more inclusions with the same inoculum than monkey kidney cell lines like Vero cells (40), and possibly LLC-MK2 cells. Overall, these data show that the transformation protocols should be further optimized for all chlamydial species (24) and that 1710S could be a potential strain to better understand plasmid maintenance dynamics in *C. pecorum*.

**Figure 3.**
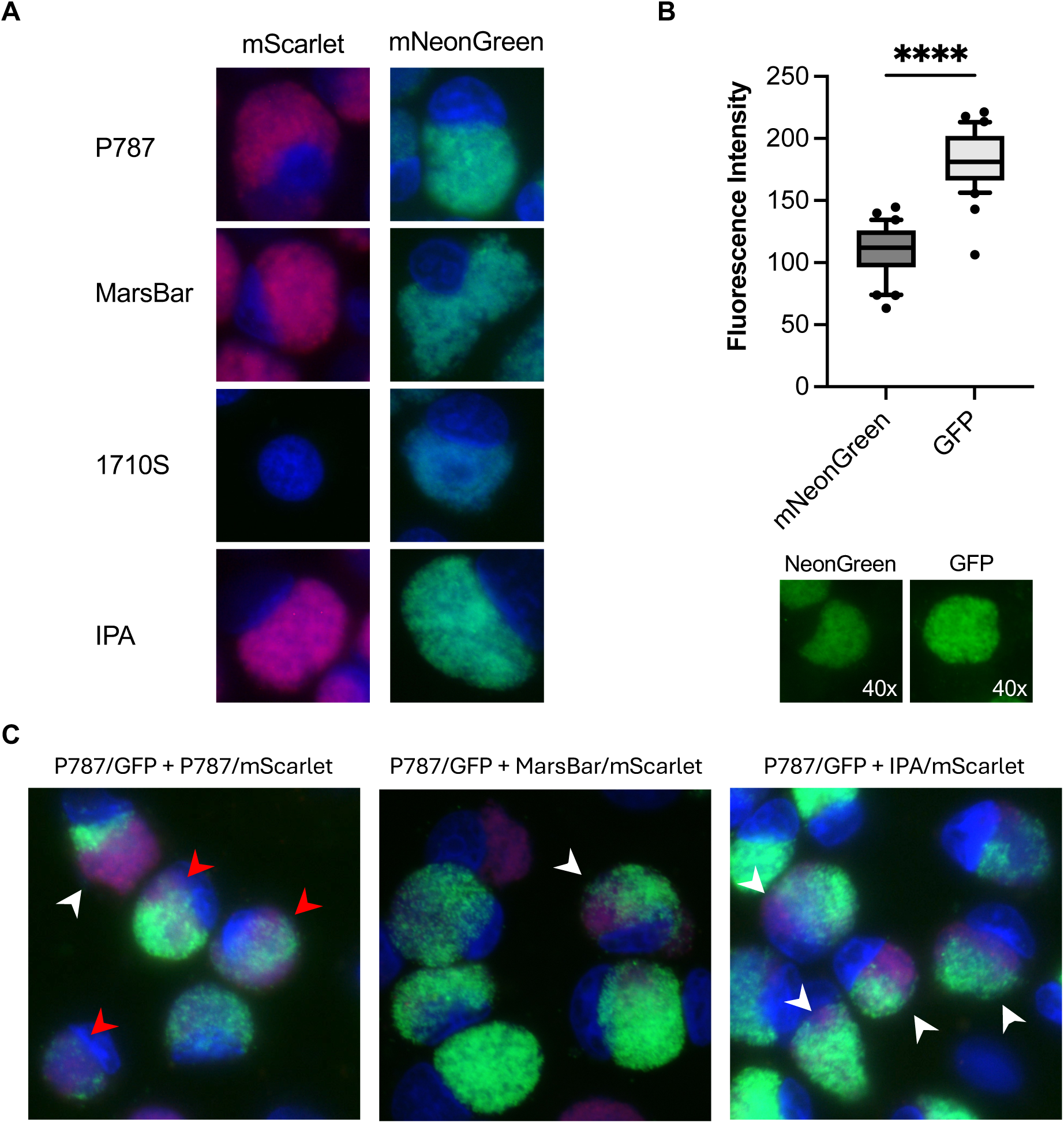
GFP-tagged inclusions yield higher fluorescence intensities than mNeonGreen-tagged chlamydiae and may be used for co-infections with mScarlet-positive chlamydiae. **A)** Shown are representative images of *C. pecorum* strains transformed with a shuttle vector containing the *C. pecorum* W73 whole-plasmid sequence and either mScarlet (left, pUC-Cpecpl-Scarlet) or mNeonGreen (right, pUC-Cpecpl-NeonGreen). Images for DAPI (blue) and mNeonGreen(green, FITC channel) or mScarlet (red, TRITC channel) were taken individually using a 60x objective, and merged. **B)** The mean fluorescence intensity of 30 *C. pecorum* inclusions tagged with NeonGreen was compared to that of GFP, using strain P787 (top). A Mann-Whitney was used for statistical analysis. Four asterisks (****) represent p-values <0.0001. Representative images for each fluorophore were taken with a 40x objective (bottom). **C)** Shown are representative images of *C. pecorum* strain P787 tagged with GFP co-cultured with mScarlet-tagged strains. Arrows indicate co-infection of the same cell with GFP- and mScarlet-positive inclusions, which are likely separate (white) or likely merged (red). Images for DAPI (blue) and GFP (green, FITC channel) or mScarlet (red, TRITC channel) were taken individually using a 60x objective, and merged.

Furthermore, we observed that the mNeonGreen green fluorescence of all *C. pecorum* strains was notably weaker than what we had seen for GFP-tagged inclusions, which stands in contrast to what we expected (64). We confirmed this observation by comparing the mean fluorescence intensity of mNeonGreen-tagged P787 inclusions to that of GFP (Figure 2B). Therefore, co-culture experiments were subsequently performed with GFP and mScarlet. We co-cultured P787/GFP with P787/mScarlet, MC/MarsBar_2018/mScarlet and IPA/mScarlet, to test for potential bleed-over between GFP and mScarlet and to investigate whether co-infection of the same cell with different strains is possible. All co-culture experiments were centrifugation-assisted (1’000 *g*, 25 °C, 1 h) with an MOI of 0.3 with clonally purified, transformed stock. Fixation was performed after 48 hours of incubation at 37 °C. We found that the bleed-over is negligible, if present. We identified the presence of different strains not only in the same cell but also possibly in the same inclusion, particularly if the same strain was used for co-infection (Figure 3C). Our results show that mScarlet and GFP are viable fluorophore options to investigate co-culture dynamics of different chlamydial strains with minimal bleed-over. Since we used 2D imaging techniques and widefield fluorescence microscopy for these initial tests, our current setup could not clearly distinguish between separate inclusions within the same cell and merged inclusions, allowing only the separation into likely separate and likely merged inclusions (Figure 3C).

## Conclusions

Overall, we were successful in transforming two veterinary chlamydial species of zoonotic importance. Similar to previous studies, we found that, while both species could be successfully transformed, the transformation rate was very low, with few to no visible transformants after the first two passages (24,65). Of note, *C. caviae* appeared to yield more transformants than *C. pecorum*. However, future optimization experiments may improve transformation success for *C. pecorum* and *C. caviae,* as previously shown for *C. suis* (24). Unfortunately, transformation of *C. abortus* with currently available protocols was not successful and requires further investigation. Moreover, to better study co-infection dynamics between different species and strains in the future, confocal microscopy or Z-stack multidimensional imaging techniques should be used to differentiate between same-cell and same-inclusion infections. Future applications of our transformants could include the study of *C. caviae* infection dissemination in guinea pigs, potential genetic exchange following intra- and interspecies co-infection, as well as the analysis of chlamydial virulence factors.

## Acknowledgments

The laboratory work was (partly) performed using the Center for Clinical Studies logistics at the Vetsuisse Faculty of the University of Zurich. We thank Barbara Sixt for donating the *C. caviae* GPIC strain, Monika Szymanska-Czerwinska for the *C. abortus* 15-70d24 strain and Kensuke Shima for helpful discussions. We also thank the laboratory team of the Institute of Veterinary Pathology for their support, especially Sandra Schneider, Theresa Pesch and Barbara Prähauser. Finally, we are grateful to Efe Altuntas and Magdalena de Arriba for their assistance in the laboratory.

## Supplementary Material

**Figure S1.**
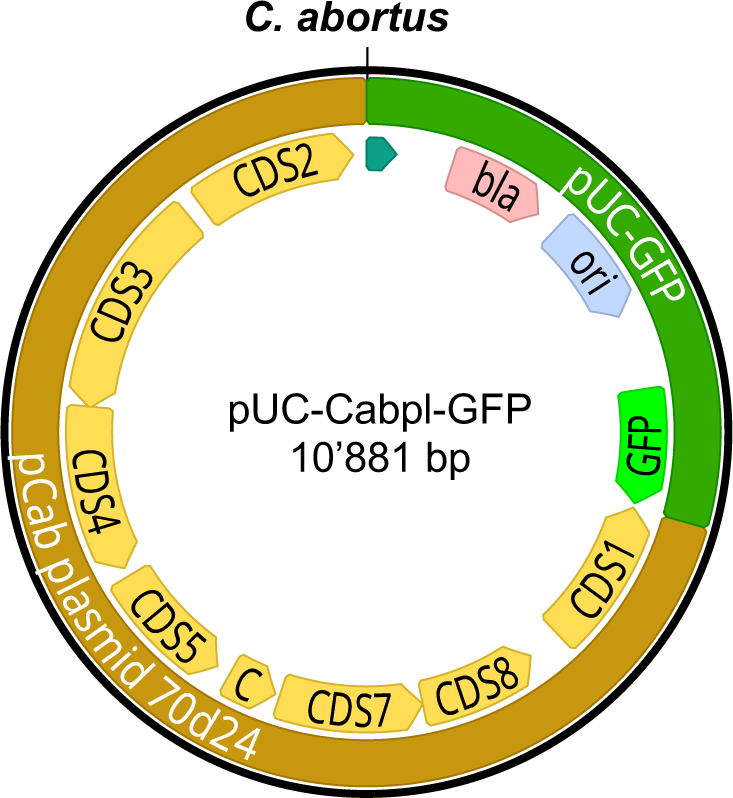
Shown is the *C. abortus* shuttle vector pUC-Cabpl-GFP comprising the native plasmid of *C. abortus* strain 15-70d24 as well as pUC-GFP, which contains a beta-lactamase (*bla,* pale red), the pUC origin of replication (*ori*, pale blue) and GFP (bright green).

**Figure S2.**
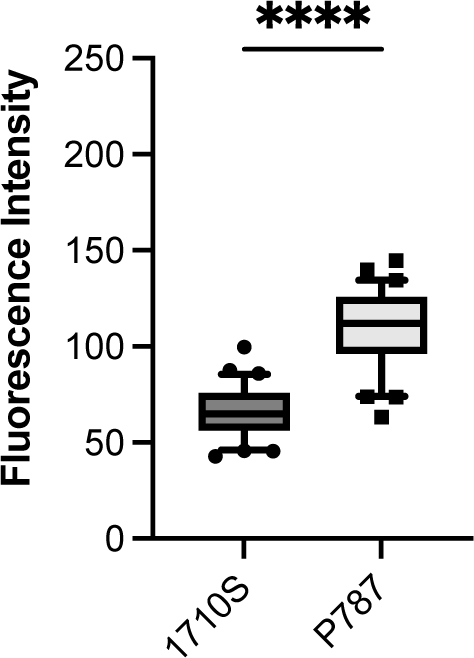
Shown is the mean fluorescence intensity of 30 inclusions of NeonGreen-tagged *C. pecorum* strain 1710S compared to P787. An unpaired t-test was used for statistical analysis. Four asterisks (****) represent p-values <0.0001.

